# A hybrid machine learning framework for functional annotation applied to mitochondrial glutathione metabolism and transport in cancers

**DOI:** 10.1101/2023.09.20.558442

**Authors:** Luke Kennedy, Jagdeep K Sandhu, Mary-Ellen Harper, Miroslava Cuperlovic-Culf

## Abstract

**Background:** Alterations of metabolism, including changes in mitochondrial and glutathione (GSH) metabolism, are a well appreciated hallmark of many cancers. Mitochondrial GSH (mGSH) transport is a poorly characterized aspect of GSH metabolism, which we investigate in the context of cancer. Existing functional annotation approaches from machine (ML) or deep learning (DL) models based only on protein sequences are unable to annotate functions in biological contexts, meaning new approaches must be developed for this task.

**Results:** We develop a flexible ML framework for functional annotation from diverse feature data. This hybrid approach leverages cancer cell line multi-omics data and other biological knowledge data as features, to uncover potential genes involved in mGSH metabolism and membrane transport in cancers. This framework achieves an average AUROC across functional annotation tasks of 0.900 and can be effectively applied to annotate a range of biological functions. For our application, classification models predict the known mGSH transporter SLC25A39 but not SLC25A40 as being highly probably related to GSH metabolism in cancers. SLC25A24 and the orphan SLC25A43 are also predicted to be associated with mGSH metabolism by this approach and structural analysis of these proteins reveal similarities in potential substrate binding regions to the binding residues of SLC25A39.

**Conclusion:** These findings have implications for a better understanding of cancer cell metabolism and novel therapeutic targets with respect to GSH metabolism through potential novel functional annotations of genes. The hybrid ML framework proposed here can be applied to other biological function classifications or multi-omics datasets to generate hypotheses in various biological contexts. Code and a tutorial for generating models and predictions in this framework are available at: https://github.com/lkenn012/mGSH_cancerClassifiers.

## Background

Glutathione (GSH) is a highly abundant tripeptide antioxidant within cells, crucial for many biological processes with its major role in regulation of reactive oxygen species (ROS). Metabolic changes are one of the hallmarks of cancer^1^, with well documented alterations in GSH metabolism foremost among them. These alterations of metabolism appear to benefit cancer cells, aiding in tumor proliferation and survival, through not yet fully understood mechanisms and interactions^2^. Alterations of GSH metabolism are essential for tumor proliferation in several cancers^3,4^ either directly where they mitigate perturbations in redox homeostasis^5,6,7^, or indirectly through ferroptosis^8^ and metabolism of chemotherapeutics^8,9^.

Central to the contributions of GSH in the metabolic characteristics of cancer cells is mitochondrial GSH (mGSH), which is found at millimolar concentrations in the organelle, at similar levels to those in the cytoplasm. Mitochondria are the primary source of ROS, largely as a byproduct of the oxidative phosphorylation system (OXPHOS), and alterations in redox balance and ROS signaling are central to the role of mitochondria in cancer cell proliferation^10^. Many GSH-utilizing enzymes with altered expression patterns in cancers are found in mitochondria, most notably glutathione peroxidase 4 (GPX4) which is a major regulator of the ferroptosis pathway. Also important are mitochondrial glutathione enzymes such as glutathione S-transferase, which conjugates and detoxifies xenobiotics and the peroxiredoxins, which lower ROS via GSH and have also been shown to promote cancer cell survival^11^.

The processes involved in the uptake of GSH into mitochondria are poorly understood. While high concentrations of GSH are present in both mitochondria and the cytosol, the synthetic enzymes are exclusively within the cytosol^12^. SLC25A10 and SLC25A11, also known as the mitochondrial dicarboxylate and oxoglutarate carriers, respectively, were initially proposed as the proteins responsible for GSH transport into mitochondria over 25 years ago^13^. However, detailed functional experiments in lipid vesicle systems in 2014 convincingly showed that no GSH transport was mediated by these two proteins^14^.

Recent evidence from two groups demonstrated mGSH transport roles for SLC25A39 and SLC25A40^15,16^. Specifically, Wang *et al.*^15^ identified the transporters through quantitative proteomics of mitochondria from GSH-depleted HeLa cells. Subsequently, Shi *et al*.^16^ leveraged CRISPR screening of gene and environment interactions to demonstrate buffering interactions between SLC25A39 and the mitochondrial iron transporter SLC25A37, revealing SLC25A39 as a candidate. The roles of these transporters in neurodegenerative diseases and cancer have recently been explored^17^. However, it remains unclear if the remaining GSH import in A39 knockouts observed in these studies is facilitated by SLC25A40, which is expressed at roughly one tenth the level of SLC25A39, or if other transport mechanisms exist, such as those for other GSH species^18,19^. Interactions affecting mGSH metabolism and transport, having secondary effects on transport may also be relevant and are even less well understood.

With recent successes of AlphaFold^20^ and RosettaFold^21^ in the elucidation of protein structure, there is a promising future for computational biology in the related problem of protein function prediction. However, competitions such as Critical Assessment of Functional Annotation^22^ have not yet found a solution for *de novo* function prediction from sequence. General function annotation models like DeepGO and DeepGOPlus^23,24^ represent major advances for the field. Along with sequence-based methods, models that leverage non-sequence features for function annotation exist^25,26^ as alternative approaches to sequence-based approaches. In addition to the general function prediction models, there are many sequence-based models designed for specific protein feature annotation such as protein-protein interactions^27^ and antibody design^28^.

Omics-based methods for functional annotation are limited. Recent work by Kunc and Kléma^29^ utilize characteristics in co-expression networks constructed from genes to predict shared KEGG pathways between genes. Similarly, Wekesa *et al.*^30^ combine differential gene expression information and knowledge of protein-protein interactions through a neighbor-voting algorithm for prediction of shared functions between yeast proteins. Finally, Wang *et al*.^31^ predict gene-gene interactions by combining co-expression features with prior biological knowledge features (e.g., subcellular localization, homology, Reactome similarity).

The approaches of Wekesa *et al.* and Wang *et al*. can be classified as hybrid ML approaches, which aim to capture “the best of both worlds” by combining the predictive powers of data-driven approaches of ML with the interpretability of theory-based models such as mechanistic or knowledge-based models. These types of models have been applied to several domains, most notably in physics-informed ML models^32^, and with respect to computational biology, hybrid modeling has been applied through diverse frameworks. Non-ML approaches have been used to uncover biological phenomena, which would otherwise be missed; a recent example integrates behavioral, transcriptomic, and network modeling to reveal the role of brain mitochondria and organization in mouse behavior^33^. Alternatively, ML models have been integrated with mechanistic models of metabolism to determine kinetic parameters and predict downstream metabolic effects^34^. P-net uses a neural network architecture based on biological knowledge of hierarchical gene-pathway-process interactions to predict prostate cancer discovery from gene features, such as methylation and copy number^35^. On the other hand, AlphaFold^20^ incorporates evolutionary information through multiple sequence alignments and physical structural constraints to inform its protein structure predictions. These hybrid models retain the predictive power of data-driven methods, often performing better than, or comparable to, standard data-driven models, while increasing the interpretability of predictions that are in-part based on biological knowledge.

With this, we sought to develop a ML framework that leverages multi-omics data from human cancer cells and existing knowledge to identify potential genes involved in mGSH transport and potential interacting metabolic processes. Several ML classifier models were developed that utilize cancer cell line encyclopedia (CCLE)^36^ transcriptomics features to predict gene ontology (GO) annotations for genes of relevant GO terms. Specifically, three classifiers were developed for annotating to: glutathione metabolic process (GO:0006749), mitochondrion (GO:0005739), and transmembrane transporter activity (GO:0022857). Additionally, we developed hybrid models for this task which include prior knowledge as features through MitoCarta^37^ scores for mitochondrial localization classification, TrSSP^38^ scores for transporter classification, or CCLE GSH & glutathione disulfide (GSSG) metabolomics data for glutathione classification. We find that Random Forest (RF) classifiers perform the best from our models tested, with hybrid models outperforming strictly transcriptomics models.

## Methods

All pre-processing, models and other computational work were conducted using in-house code written in Python (Python Software Foundation. Python Language Reference, version 3.8. Available at www.python.org). All algorithms and methods used in ML model building and training were implemented via Python’s scikit-learn package^39^ unless stated otherwise. Figures for evaluations of classifier models were produced via the matplotlib and seaborn libraries^40,41^; GO enrichment plots were produced using ShinyGO 0.77^42^; and all other images were created with BioRender.com.

### Data collection & pre-processing

#### Metabolomic & transcriptomic CCLE data

The Broad Institute’s Cancer Cell Line Encyclopedia (CCLE) provides multi-omics data across over 1,000 human cancer cell lines (CCLs). CCLE transcriptomics, and metabolomics data were used in building out models^36,43^. Raw transcriptomics and metabolomics data were downloaded from broadinstitute.org (available at the time of writing via depmap.org/portal/download/).

Data were cleaned and imputed by removing all CCLs and transcripts/metabolites with >30% missing values or standard deviation values of 0 across samples. Remaining missing values were imputed using the k-nearest neighbors (k-NN) imputation algorithm (using Euclidean distance with k=5 selected to minimize impact of the data structure on the missing data imputation^44^). For metabolomics features, Spearman correlations were generated across common CCLs between GSH, GSSG metabolomics data and transcripts in CCLE transcriptomics data. Non-significant correlations (p-value ≥ 0.05, Student’s t-test with 2 degrees of freedom) were set to 0. Spearman rho values were normalized by Fisher transformation^45^.

Feature space reduction while selecting major variances in CCLE transcriptomics was performed using principal component analysis (PCA). These principal components (PCs) were used as features in classifier models (further described in Model development and feature selection process).

#### Knowledge-based features

Mitochondrial localization scores for genes were based on MitoCarta 3.0^37^ (broadinstitute.org/mitocarta). MitoCarta scores are determined through a naive-Bayes model which combines features from several independent domains from features used in our analysis, such as homology, sequence domain, and tandem mass spectrometry of purified mitochondria. CCLE genes were assigned MitoCarta scores according to these data, with 1 indicating mitochondrial localization, and 0 indicating non-mitochondrial localization. For genes that are not assigned scores in MitoCarta, a value of −1 is assigned.

Transporter activity was based on TrSSP^38^ values, obtained from Zhao lab webpage (www.zhaolab.org/TrSSP). TrSSP utilizes support vector machine models to predict membrane transport proteins and their substrate classes using primary protein sequence features along with position-specific scoring matrices to predict transport function and specificity. Like MitoCarta scores, TrSSP scores were incorporated into the model as classifications for positive, negative, or no prediction by TrSSP (1, 0, −1, respectively).

For use as features in our classifiers, these knowledge-based categorical scores were transformed to Boolean features using one-hot encoding.

### mGSH transporter classifier models

#### Training and test gene sets selection

For each classifier, training and test genes were selected based on existing knowledge of gene functions. Gene ontology^46^ terms were used to determine genes related to mGSH metabolism: glutathione metabolic process GO:0006749 (65 genes), mitochondrion GO:0005739 (1685 genes), transmembrane transporter activity GO:0022857 (1154 genes). These terms cover all classes of ontology terms: biological process, cellular component, and molecular function, respectively. Annotated genes for each term were identified via AmiGO 2^47^ and those genes annotated based on only low confidence computational or inferred evidence were removed. This left 40 genes for GO:0006749, 781 genes for GO:0005739, and 700 genes for GO:0022857 annotated based on experimental evidence to create the final list of positive class genes for the classifier models. We found that there is little overlap among these gene sets; primarily, common genes are found between mitochondrial and transmembrane transporter activity genes (72 genes), with 5 genes overlapping the mitochondrial and glutathione metabolic process sets and only one gene found in both transmembrane transporter activity and glutathione metabolic process genes. There are no genes annotated by all three GO terms. AmiGO annotations based on sequence similarities were retained along with the experimental AmiGO annotations used for the other GO annotations to generate the 40 genes used in the final positive gene set for glutathione metabolic process term. These annotations were retained due to the small number of genes annotated by experimental evidence for the GO term, thus creating a larger positive gene set for model training. bioDBnet^48^ was used to link gene symbols from AmiGO to Ensembl gene IDs found in the CCLE data.

The negative classification set of genes was generated by randomly undersampling from genes that were not included in the positive set based on GO annotations. By undersampling, positive and negative genes (samples) for model training and testing were balanced. However, training a model through this process is biased by the specific set of randomly selected negative class genes in the training set. To ensure robustness of the models, 100 iterations of random sampling were performed to create different sets of samples for model training. Final predictions and evaluations are average results over training and testing iterations on the 100 different data sets.

### Model development and feature selection process

To determine the best methodology for our task, several common ML algorithms and feature sets were tested. Random Forest (RF), Decision Tree (DT), Naive Bayes (NB), and Support Vector Machine (SVM) were applied for classification tasks of mitochondrial, transmembrane transporter activity, and GSH metabolic process GO terms. Models were tested with several sets of features that combine experimental features from CCLE data with knowledge-based features, or feature sets containing only experimental features. Experimental feature sets include the first 5, 14, 30, or 50 CCLE transcriptomics PCs which capture 93%, 95%, 97%, or 98% of the variance, respectively. For classifying the GSH metabolic process GO term, Spearman correlations values between gene transcriptomics and GSH or GSSG metabolomics were included as features as well. Features from existing biological knowledge were included through MitoCarta scores for mitochondrial localization in the mitochondrial classification task, and TrSSP prediction scores for membrane transport proteins in the transporter classification task.

For comparisons of feature sets in models, either metabolomics or knowledge features were replaced with additional PC features to create the transcriptomics-only models with an equivalent number of features (i.e., a mitochondrial classifier with the first 5 PCs and MitoCarta scores as features is comparable to a transcriptomics-only classifier using the first 6 PCs as features).

### Model evaluation

Model performance was analyzed from predicted class probabilities using common evaluation metrics including receiver operating characteristic (ROC) and precision-recall curves (PRC). These methods provide a balanced approach to evaluating classifier performance, which considers the effects of true positives (TPs), true negative (TNs), false positives (FPs), and false negatives (FNs). Models were evaluated and selected based on values for the areas under ROC (AUROC) and PRC (AUPRC), and Matthew’s correlation coefficient (MCC)^49,50^. These evaluation metrics were calculated across all test-set predictions from five-fold cross-validations of randomly sampled training/test sets. Reported metrics are calculated as average values across 100 iterations of training/testing using the different gene sets.

### Protein structural analysis

#### SLC25 structures

Protein structures are obtained from the AlphaFold^20^ structure database (alphafold.ebi.ac.uk) for use in structural analyses. Additionally, homology models from SWISS-MODEL^51^ for both SLC25A39 and A40 are included in our analyses. Homology based structures are modeled on the crystal structure of the bovine ADP/ATP carrier (PDB:OKC1)^52^ and cryo-electron microscopy determined structure of human UCP1 (PDB:8HBV)^53^ for A39 and A40, respectively.

#### Protein structure alignments

Comparisons of 3D protein structures were conducted using sequence-independent alignment through the TM-align algorithm^54^ and quantified by the corresponding TM-scores, which quantifies similarities through similarities in protein structure topology. This method performs comparably to other common alignment methods and does not consider sequence similarity for alignment, which could skew the alignments due to conserved sequences within the SLC25 family. Alignments were implemented using the TMalign module for PyMOL^55^ and predicted mGSH transporter structures were compared to known mGSH transporters (SLC25A39, A40).

The CAVER PyMOL plugin version 3.0.3^56^ was used to identify tunnel residues for each SLC25 structure. Tunnels were identified by initializing the tunneling at the base or innermost point of the transporters tunnel with maximum starting point distance and desired radius of 3Å for starting point optimization. From this base, the tunneling algorithm probed outward to generate an interior tunnel channel. CAVER default parameters were specified for the tunneling algorithm, using 12 approximating balls with minimum probe radius of 1.5Å, shell depth of 20Å, shell radius of 7Å (or 10Å for structures where a smaller radius fails to fill the tunnel space), and a clustering threshold of 3.5. Default parameters were used for computation memory and speed, and were adjusted as described, if necessary, for finding transporter tunnels. Relevant tunnels were identified from the top-ranked tunnels identified by CAVER as those tunnels which traversed and filled the length of the proteins’ tunnel without escaping to the protein surface through a side gap or by “overflowing” the interior tunnel.

In the same fashion as the full structure alignments, analysis of the tunnel regions for the SLC25 family were performed by aligning the tunnel-interacting residues identified from CAVER using TM-align.

## Results

### Classifier models for mGSH transporters in cancer

To identify top mGSH transporter candidates, three classifier models were developed to classify genes based on our desired candidate characteristics: mitochondrial localization, association with GSH metabolism, and transmembrane transport function (see **Fig.** 1.a. for overview and **Methods** for details). Each model uses features from CCLE transcriptomics data along with one or more features from other sources; GSH and GSSG metabolomics correlations for GSH metabolic process classification, MitoCarta scores for mitochondrial localization classification, and TrSSP scores for transmembrane transporter activity classification (**Fig.** 1. b.).

**Figure 1:**
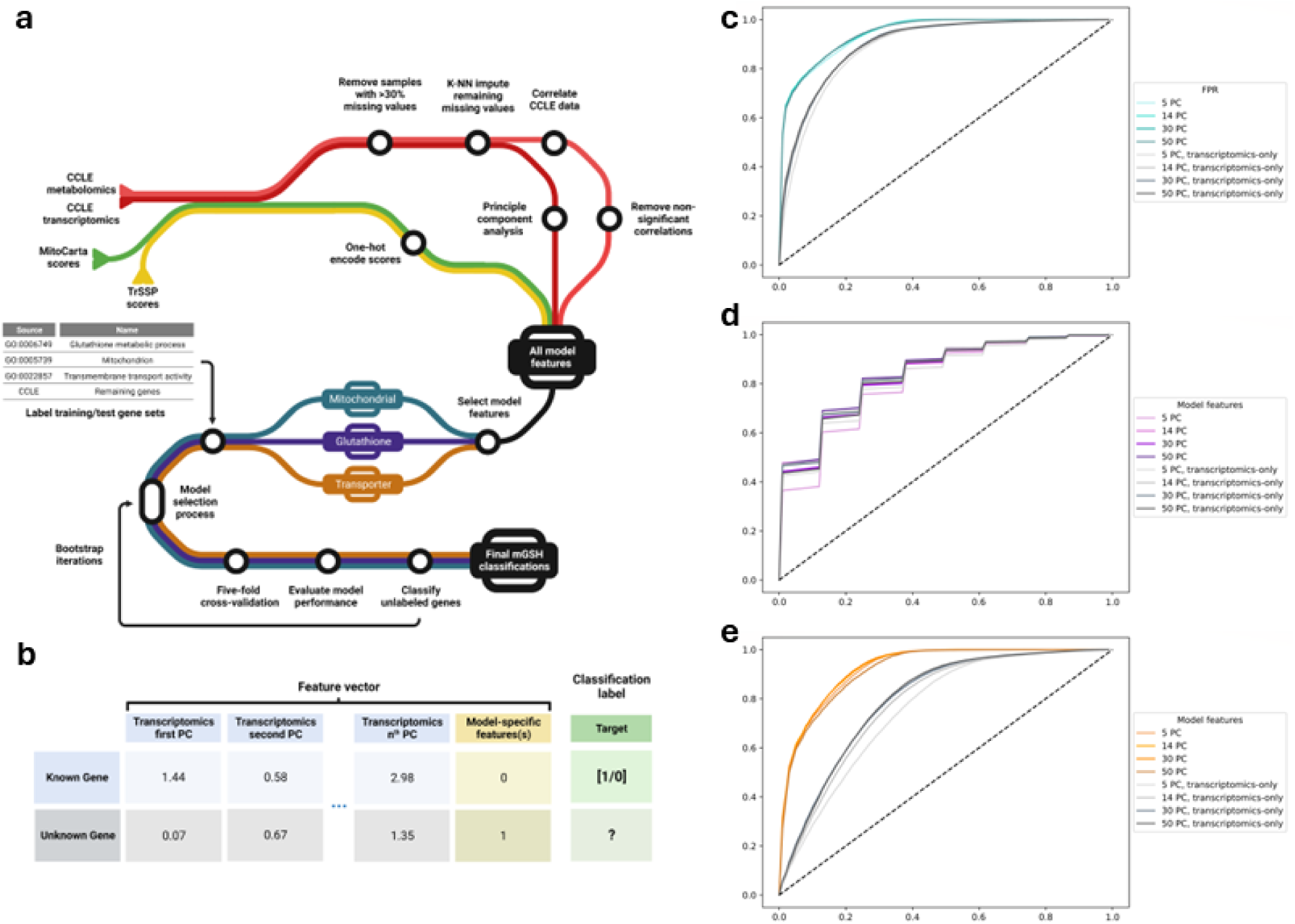
Overview of mGSH transporter classifier model design and evaluation. (**a**) Schematic of the workflow for ML model development (inspired by designs by Fellows Yates *et al.*^85^). In this diagram, lines represent the flow of information from the initially collected raw data through several transformation steps to the processed model features, and then through the ML training, evaluation, and selection steps to obtain the final classification of our unknown genes. (**b**) Example feature vectors assigned to genes in this framework, where transcriptomic features are PCs and model-specific features are MitoCarta scores, for example. ROC curves are presented for mitochondrial (**c**), transporter (**d**) and glutathione (**e**) classifiers with various feature sets. Models which incorporate transcriptomics and other feature data are colored while transcriptomics-only models are gray. Differently shaded curves represent models with various numbers of transcriptomics features (range: 5-50).

Our classifiers performed well in the tasks of mitochondrial, GSH, and transporter annotation classification across our evaluation metrics, as shown in **Table 1**. Across all models and feature selections, mean five-fold cross-validations shows AUROC is 0.820, and RF classifiers being the best performer with mean AUROC values of 0.900. Classifier performance varied across our three classifier types, with mitochondrial classifiers performing the best and glutathione classifiers performing the worst, on average, which may be attributed to two factors. First, the set of annotated genes related to GSH metabolism is much smaller than the gene sets for the other classifiers (∼40 genes vs. ∼700 genes), thus providing a smaller training set for these models. Additionally, the GSH classifiers rely solely on CCLE multi-omics data unlike the other two classifiers which also include data from existing knowledge-sources.

**Table 1.** Average performance of classifiers in mGSH transporter classification. . Scores are mean values across each classifier type and all feature sets which include transcriptomic and other data. Individual scores are calculated as averages across bootstrap iterations. The highest scoring model for each evaluation metric is bolded.

| Algorithm | Metric |  |  |
| --- | --- | --- | --- |
|  | AUROC | AUPRC | MCC |
| Decision tree | 0.7663 | 0.8270 | 0.5371 |
|  | 0.6896<sup>†</sup> | 0.7692<sup>†</sup> | 0.3840<sup>†</sup> |
| Naive-Bayes | 0.8553 | 0.8422 | 0.3517* |
|  | 0.7433<sup>†</sup> | 0.7334<sup>†</sup> | 0.2028<sup>†</sup>* |
| Random forest | **0.9003** | **0.8941** | **0.6349** |
|  | **0.8380<sup>†</sup>** | **0.8156<sup>†</sup>** | **0.5437<sup>†</sup>** |
| Support vector machine | 0.7614 | 0.7558 | 0.3487 |
|  | 0.7130<sup>†</sup> | 0.7250<sup>†</sup> | 0.3125<sup>†</sup> |

<sup>†</sup> scores from models with transcriptomics-only features.
\* low MCC values relative to other evaluation metrics due to low positive class accuracy.

Along with testing several ML algorithms in our classifiers, multiple feature sets were explored to determine the best selection for our task. Specifically, variations of the number of top PCs as linear combinations of transcriptomic features (described in Methods) from 5-50, were tested. Increasing the number of these features in our models did not notably improve performance by our metrics (**Supp. Fig.** 1). Based on these results, a small number of the first several PCs are sufficient for effective classification. However, specific gene classification probabilities are variable across models with different numbers of PC features used. Since performance is similar across these models, rather than selecting a specific model, we used mean gene classification probabilities with standard error (SE) calculated from models with different numbers of PC features (5, 14, 30, 50) for final classification results.

### Comparison of hybrid models and transcriptomics-based models

Consistently across all tested models, classifiers that incorporate both experimental data from CCLE transcriptomics data and data from other knowledge bases as features outperform those models with an equivalent number of experimental data features alone (**Fig**. 1.c.,d., **Table 1**). Although no knowledge component exists for the GSH classifiers, transcriptomics-based models were compared to models which incorporate GSH and GSSG metabolomics data from the CCLE along with the CCLE transcriptomics data for features. Interestingly, the additional data source does not seem to similarly improve performance in the GSH models, with performance of transcriptomics-only GSH classifiers comparable to GSH classifiers using both transcriptomic and metabolomic features (**Fig.** 1.e.).

### Comparison of hybrid models to knowledge-based models

Like the previous analysis, we sought to evaluate the effects of adding transcriptomics information to the existing predictive models from the literature. Classifications from our mitochondrial and transporter models were compared to those of MitoCarta and TrSSP. The combined transcriptomic and MitoCarta/TrSSP classifiers appear to only show slight differences (**Fig.** 2.a.,b.). The mitochondrial classifier identifies slightly more mitochondrial genes than MitoCarta alone, however, there are more false positives (FPs) in this classifier (at a classification threshold of 0.75).

**Figure 2:**
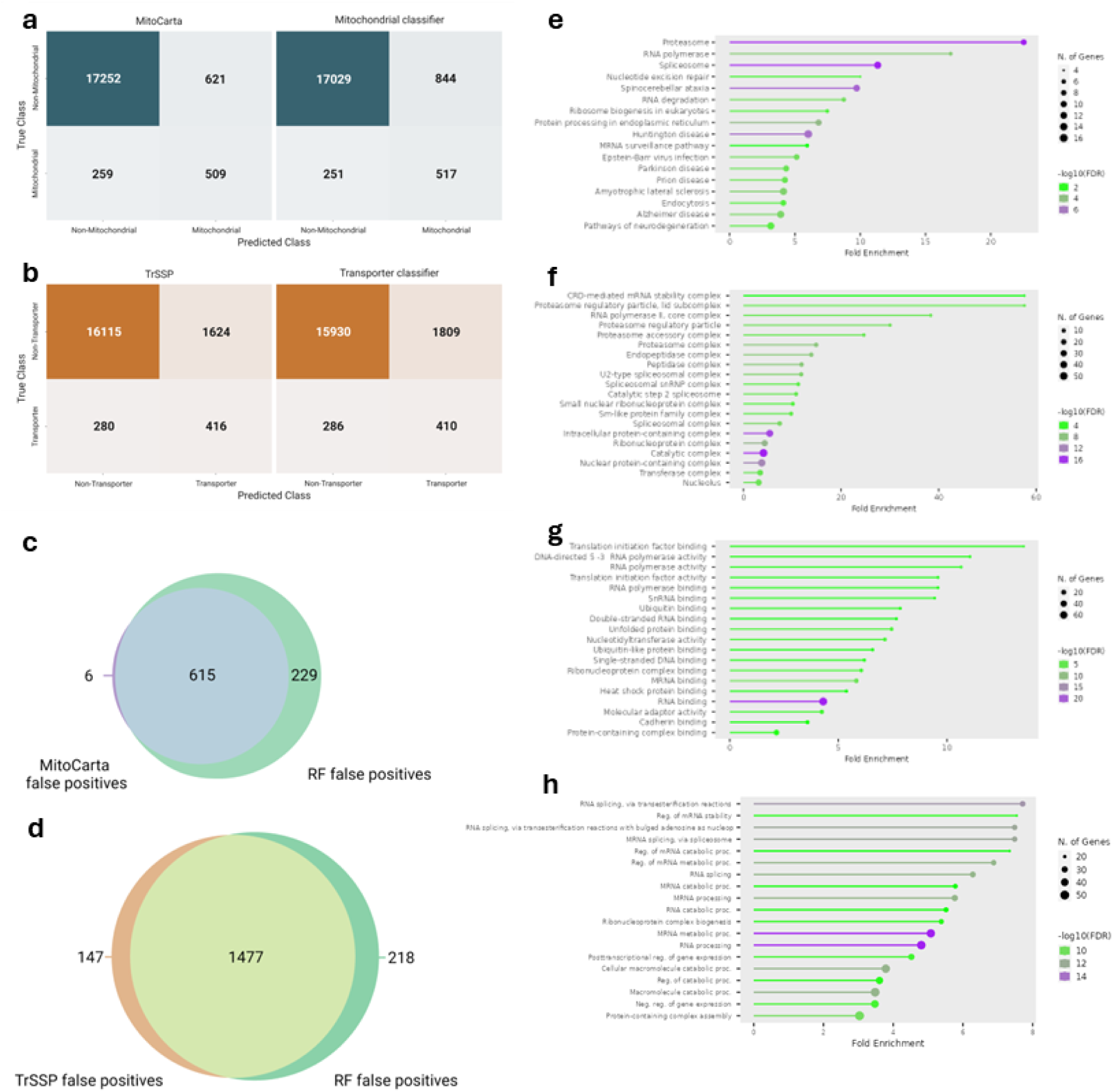
Confusion matrices of existing mitochondrial and transporter models (left) and models produced here (right). (**a**) MitoCarta compared to our mitochondrial RF classifier. (**b**) TrSSP compared to our transporter RF classifier. Matrices are colored based on the number of samples contained within each quadrant. A classification threshold of 0.75 was used for the mitochondrial and transporter classifiers as it provided the best balance of maximizing true positives and minimizing false positives. (**c**) Venn diagram of false positive mitochondrial genes predicted by MitoCarta 3.0 and by the mitochondrial RF classifier produced here. (**d**) Venn diagram of false positive transporter genes predicted by TrSSP and by the transporter RF classifier produced here. (**e**-**h**) GO enrichment analysis of genes predicted to localize to mitochondria by RF classifiers with no existing evidence in localization databases. Enrichment plots of false positive mitochondrial genes for KEGG pathways (**e**) cellular component (**f**), molecular function (**g**), and biological process (**h**) GO terms.

### False positive mitochondrial and transporter genes

Interestingly, our classifiers predict more false positives compared to the results of MitoCarta and TrSSP, suggesting potential for identifying novel functions. To assess this, mitochondrial FP genes were compared to other sources of mitochondrial genes beyond the high confidence GO annotations used for classifier training. Low confidence GO annotations that were originally removed from the training set, Human Protein Atlas subcellular localization annotations from histology images^57^, Swiss-Prot annotations^58^, and integrated mitochondrial protein index (IMPI) annotations based on MitoMiner^59^ were used for other sources of localization evidence. Of the 844 FP genes predicted by the mitochondrial RF classifier, 615 were predicted by both MitoCarta and RF, and 229 were predicted by RF only (**Fig.** 2.c.). 206 of these RF FP genes have no evidence in the other datasets, while the remaining have evidence in at least one source (**Supp. Data**). Enrichment analysis revealed that the 206 unannotated FPs without existing evidence are significantly enriched in KEGG pathways (**Fig.** 2.e.) and GO terms (**Fig.** 2.f.-h.) that are associated with degradation pathways, such as the proteasome, RNA stability, degradation and processing pathways, and are furthermore highly associated with one another based on high confidence evidence through STRING^60^ (**Supp. Fig.** 4). The proteasome system is closely linked to mitochondrial remodeling in several physiological and disease states, including cancer^61,62^.

Comparisons of FPs in TrSSP and our transporter RF classifier find less overlap between the two (**Fig.** 2.d.). 147 FPs are exclusive to TrSSP, 218 are exclusive to the RF classifier, and 1477 are common to both models. In the same fashion as mitochondrial genes, other evidence sources beyond the high confidence GO term annotations were compared to the FPs predicted by transporter models. Low confidence AMiGO annotations, Transporter Classifier Data Base annotations^63^, and the SLCAtlas^64^ were used as alternate evidence sources. Swiss-Prot annotations were excluded as these were used in TrSSP model training. For both TrSSP and our RF classifier, only a small number of FP genes have evidence from other sources (2 for TrSSP, and 8 for the RF classifier, **Supp. Data**). Unlike the mitochondrial FPs by our RF classifier, transporter FP genes are not extensively enriched in specific GO terms (**Supp. Fig.** 5), and no enrichment of KEGG pathways is observed. The most apparent enrichment is in biological process terms related to cell-cell interactions, such as cell adhesion, cell junctions, and synapses (**Supp. Fig.** 5.c.).

### mGSH transporter candidates

The results of classifiers identify several potential mGSH transporters. Within the SLC25 family, the known transporter SLC25A39 is among the most probable GSH-related genes (**Table 2**). Surprisingly, SLC25A40, the homolog of SLC25A39, has a very low GSH probability by the RF classifier models and is not predicted to be related to GSH. Amongst the top candidates are several well characterized proteins such as SLC25A1, SLC25A10, and A11 as well as some orphan members like SLC25A43 and A50. To understand the possible roles of these members in GSH transport, existing evidence was considered and described below.

**Table 2:** Most probable GSH transporting SLC25 proteins by mean GSH RF classifier probability. . GSH probabilities are mean probabilities with standard error for RF classifiers over different feature sets. The known GSH transporters SLC25A39 and SLC25A40 are in bold.

| Rank | Gene Symbol | GSH probability |
| --- | --- | --- |
| 1 | SLC25A1 | 0.7891 (0.0173) |
| 2 | SLC25A10 | 0.7800 (0.0256) |
| 3 | SLC25A13 | 0.7737 (0.0156) |
| 1 | SLC25A1 | 0.7891 (0.0173) |
| 2 | SLC25A10 | 0.7800 (0.0256) |
| 3 | SLC25A13 | 0.7737 (0.0156) |
| **4** | **SLC25A39** | **0.7716 (0.0230)** |
| 5 | SLC25A50 | 0.7620 (0.0222) |
| 6 | SLC25A43 | 0.7603 (0.0318) |
| 7 | SLC25A24 | 0.7577 (0.0345) |
| 8 | SLC25A37 | 0.7494 (0.0260) |
| 9 | SLC25A3 | 0.7410 (0.0489) |
| 10 | SLC25A11 | 0.7322 (0.0074) |
| **37** | **SLC25A40** | **0.4702 (0.0152)** |

### Validation of annotation framework

Several other metabolite classifier models were developed for the mGSH transporter candidates identified by this approach. Like the GSH classifiers, RF models with 5-50 CCLE transcriptomic PCs and corresponding metabolomic correlations as features were trained for the annotation of genes related to 2-oxoglutarate, glutamate, and carnitine.

Following the procedure for the GSH classification task, positively classified genes were identified from GO terms for these metabolites to generate training and test gene sets for classifiers. 2-oxoglutarate, glutamate, and carnitine were selected for this evaluation because, out of metabolites found in the CCLE metabolomics set, they have a relatively larger number of annotated genes, making classifier training possible. Additionally, there is evidence for transport by each for one or more SLC25 proteins, which allows for some measure of classifier validation. For each of these classifiers, performance was comparable to the GSH classifier by our evaluation metrics (**Supp.** **Table** 1, **Table 3**).

**Table 3:**
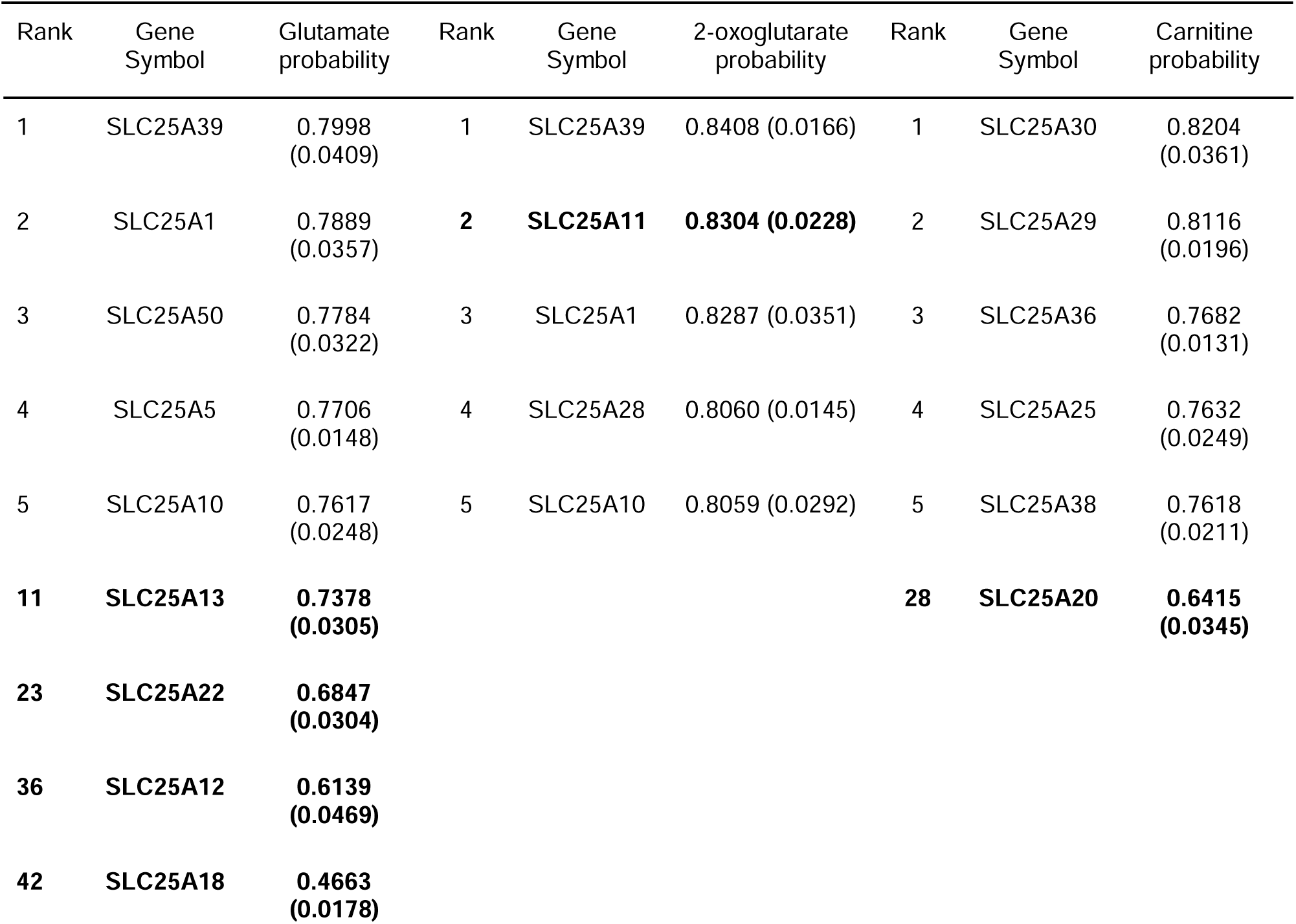
Top SLC25 candidates for non-GSH metabolites by RF classifiers. . Candidates ranked by mean RF classifier probabilities (5-50 PC features) with standard error. Known metabolite transporters are bolded.

| Rank | Gene Symbol | Glutamate probability | Rank | Gene Symbol | 2-oxoglutarate probability | Rank | Gene Symbol | Carnitine probability |
| --- | --- | --- | --- | --- | --- | --- | --- | --- |
| 1 | SLC25A39 | 0.7998 (0.0409) | 1 | SLC25A39 | 0.8408 (0.0166) | 1 | SLC25A30 | 0.8204 (0.0361) |
| 2 | SLC25A1 | 0.7889 (0.0357) | **2** | **SLC25A11** | **0.8304 (0.0228)** | 2 | SLC25A29 | 0.8116 (0.0196) |
| 3 | SLC25A50 | 0.7784 (0.0322) | 3 | SLC25A1 | 0.8287 (0.0351) | 3 | SLC25A36 | 0.7682 (0.0131) |
| 4 | SLC25A5 | 0.7706 (0.0148) | 4 | SLC25A28 | 0.8060 (0.0145) | 4 | SLC25A25 | 0.7632 (0.0249) |
| 5 | SLC25A10 | 0.7617 (0.0248) | 5 | SLC25A10 | 0.8059 (0.0292) | 5 | SLC25A38 | 0.7618 (0.0211) |
| **11** | **SLC25A13** | **0.7378 (0.0305)** |  |  |  | **28** | **SLC25A20** | **0.6415 (0.0345)** |
| **23** | **SLC25A22** | **0.6847 (0.0304)** |  |  |  |  |  |  |
| **36** | **SLC25A12** | **0.6139 (0.0469)** |  |  |  |  |  |  |
| **42** | **SLC25A18** | **0.4663 (0.0178)** |  |  |  |  |  |  |

### Non-SLC25 mGSH-related proteins

Alongside the SLC25 family, the remaining genes in our dataset were considered for roles in mGSH transport. Top candidates were identified by their mean GSH probability across RF classifiers. Additionally, the mean mitochondrial and transporter classifier scores were considered. To identify high confidence candidates, only genes with probabilities greater than 0.80 across all classifications are selected (i.e., genes already annotated with one of our GO terms are said to have probabilities of 1). This thresholding removes 99.95% of the available 49309 genes in the datasets, leaving only 27 genes that meet our criteria (**Supp.** **Table** 2). Of these, 4 genes are already annotated with the glutathione metabolic process GO term, 13 are annotated with the mitochondrion GO term, and 5 with the transmembrane transporter activity term.

### Structural comparisons

To further examine the SLC25 proteins most probably relevant to GSH transport, we next conducted structural analyses. Experimental determinations of protein structures for the SLC25 family are currently lacking. The first complete human SLC25 structure was recently determined for UCP1 by cryo-electron microscopy^53^. Beyond this, structures for orthologs in model species have been determined through x-ray crystallography for the bovine^54^ and *Thermothelomyces thermophila*^65^ ADP/ATP carriers (SLC25A4), and by NMR molecular fragment searching for the yeast UCP2 (SLC25A8)^66^. Predicted protein structures must be used for comparison of family members with unknown structures. Particularly, we were interested in structural comparisons between our candidates with the GSH transporters SLC25A39, and SLC25A40 to reveal any similarities in structure and potential function. Pairwise structural alignments were determined using TM-align (see Methods) for full 3D structures (“global” alignments) or tunnel-interacting residues within structures (“local” alignments). The residues for local alignments were identified via CAVER^56^ (**Supp. Fig.** 6). Rather than considering the entire transporter structure, of more relevance to transporter function and specificity is the interior tunnel of the protein surface, along which substrates interact and move across the IMM^65,67^. By aligning only potentially substrate binding residues we get a more relevant analysis of the SLC25 family with respect to substrate binding as these structures are expected to be more variable and better indicators of shared transport activities.

Pairwise global alignments showed high similarities (TM-scores close to 1) across the transporter family (**Supp. Data**). This is expected due to common structural domains across the family, namely the six transmembrane helices. TM-scores for local alignments of tunnel residues are much lower relative to the global alignment scores (mean scores excluding self-alignments 0.376 vs. 0.701) (**Supp. Fig.** 7) and more variable (coefficient of variation of 0.350 vs. 0.172 for global alignments), indicating the diversity of transport tunnels and functions of the SLC25 proteins. **Figure** 3.a. details the TM-scores for local pairwise alignments between top mGSH candidate SLC25s as well as SLC25A39 and SLC25A40.

**Figure 3:**
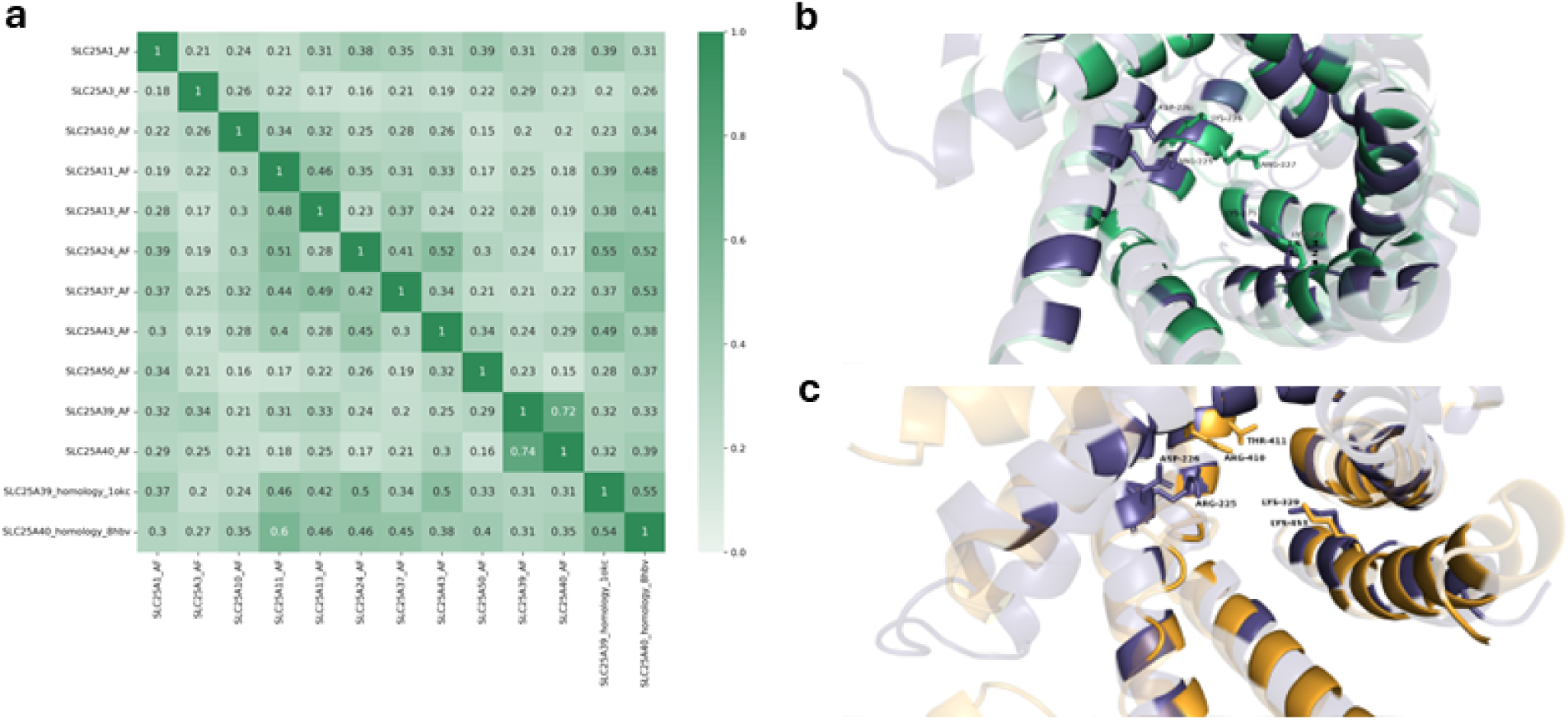
Structural alignment by TM-align of CAVER-identified transporter tunnels in SLC25 family members predicted to be associated with glutathione metabolic processes. (**a**) Heatmap of TM-scores for pairwise alignments of tunnel-interacting regions identified by CAVER for candidate SLC25 members. Values are TM-scores normalized to the sequence length of the column protein. Highest scoring alignments for each column are bolded. (**b**) & (**c**) Pymol visualization of transporter tunnel alignments for SLC25A39 (dark purple) and most similar SLC25 candidates by TM-score, SLC25A43 (green) and SLC25A24 (orange). Indicated residues are those identified as relevant for GSH transport by SLC25A39 and corresponding residues in SLC25A43 and SLC25A24. Protein tunnels visualized from the intermembrane space with tunnel-interacting residues identified by CAVER highlighted and non-tunnel residues faded.

Amongst these results, relatively high scores between SLC25A24, SLC25A43, and SLC25A39 indicate similarities in transporter tunnels and possible overlapping transport functions of the proteins. Shi *et al*.^16^ identify R225, D226, and K329 as important residues for GSH transport in SLC25A39. Based on local structural alignments of tunnel regions with SLC25A43 and SLC25A24 (**Fig.** 3.b. & c.), the position of the K329 residue is conserved by K275 in SLC25A43 and by K453 in SLC25A24. However, the R225, D226 motif of SLC25A39 appears less conserved in the aligned structures. In the SLC25A43 alignment, residues G173 and A174 are in closest proximity to the A39 binding residues, and these non-polar residues are not identified by CAVER as tunnel-interacting residues. Interestingly, the sidechains of SLC25A43 residues K226 and R227 appear near R225 and D226 sidechains in SLC25A39, where the positively charged K226 replaces the negative charge of D226 in SLC25A39. Like SLC25A43, the closest aligned residues of SLC25A24 to the GSH binding residues of SLC25A39 are the non-polar G353 and I354, which do not interact with the transporter tunnel. Where K226 and R227 are found in SLC25A43, instead there is R410 and T411 in SLC25A24.

### Comparison to DeepGOPlus

For a comparison to the models developed here, genes from our GO terms of interest were annotated by DeepGOPlus. DeepGOPlus^24^ is a general function annotation deep learning model based on protein amino acid sequences. The model is constructed as a convolutional neural network (CNN) in which protein sequences are taken as inputs and the predicted GO term annotation and probabilities are returned. Gene ontology terms with predicted annotation probabilities were obtained through the DeepGOWeb API^68^ (Version 1.0.13) using FASTA sequences obtained from canonical Ensembl transcript IDs for the genes under each ontology term. All 39 high confidence glutathione metabolic process annotations, and 150 randomly selected genes from each other ontology term were annotated using DeepGOWeb. The true positive rates (TPRs) or sensitivity measures of annotations were determined for model comparison (**Fig.** 4, **Supp. Table** 3).

**Figure 4:**
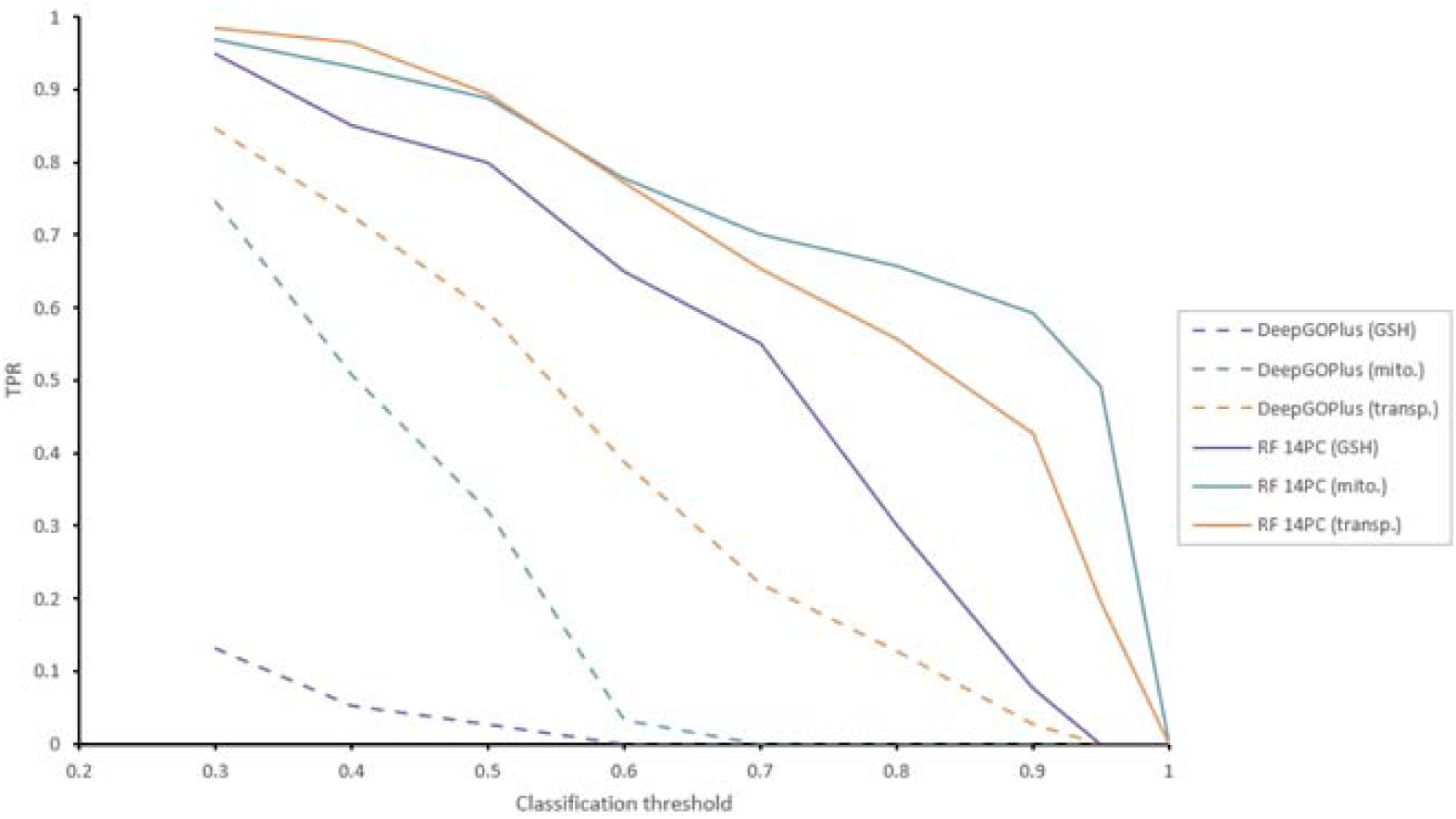
Evaluation of gene ontology annotation predictions by DeepGOPlus (dashed) and RF classifier models (solid). Sensitivity, or TPR, are calculated at various classification thresholds.

Our classifiers perform better than DeepGOPlus for annotating all GO terms of interest. This is most evident for the set of genes involved in glutathione metabolism, where DeepGOPlus fails to annotate this set of genes with an average TPR of only 0.073 over classification thresholds compared to 0.867 for the RF classifier. DeepGOPlus can reasonably classify mitochondrial localization and transporter activity annotations with mean TPRs of 0.525 and 0.722, respectively.

## Discussion

The importance of mitochondria and GSH in cancers is widely appreciated. Mitochondria have a major role in metabolic shifts, one of the hallmarks of cancer, while mGSH is important in metabolism, ROS processing, post-translational protein modifications and tumor drug resistance. An improved understanding of mitochondrial transport mechanisms in cancer cells, including GSH transport mechanisms, is important for advancing the cancer biology field. To this end we developed a hybrid ML framework that utilizes multi-omics data and existing knowledge for annotating genes with GO terms of interest. We applied this model to the problem of mitochondrial GSH metabolism, specifically focusing on the aspect of transport. We propose several potential candidates that are predicted to be related to, and possibly modulated by, mGSH transport that are identified by this model to guide future experimentation in this area of research.

Random forest classifiers were developed here for the annotation of mGSH transporter characteristics: mitochondrial localization, relation to GSH metabolism, and transporter function. Models combine CCLE transcriptomics and metabolomics data for annotation of genes related to GSH metabolism, and knowledge from existing predictive models for mitochondrial localization and transporter activity annotation. This framework resulted in models with high performance (mean RF AUROC of 0.900) across each classification task.

For comparison, we attempted similar classifications using DeepGOPlus^24,68^ for annotation of our terms of interest. This general GO annotation model performed worse, as measured by sensitivity in correctly classifying annotated genes, than our framework This is most evident in GSH metabolic process classification, likely due to the relatively small set of annotated genes for this term which is underrepresented during training in the DeepGOPlus framework, indicating a weakness of these general frameworks and a strength of our approach. Furthermore, as a sequence-based approach, DeepGOPlus cannot be applied to disease contexts as we do with our proposed method, in this case using cancer cell transcriptomics and metabolomics data to focus analysis on a cancer context.

As a further validation of our methodology, several other classifiers using our framework were developed and tested for annotating other GO terms. Identical RF models to the GSH classifier were developed which instead use genes annotated with GO terms for glutamate, 2-oxoglutarate, and carnitine metabolic process for training. These models show similar performance to the GSH classifier despite a small number of annotated genes for training, indicating the robustness of this framework for annotating a variety of biological functions. Furthermore, with respect to our interest in transport, these classifiers correctly identify many previously known SLC25 transporters as their most probable candidates.

Considering the SLC25 family of mitochondrial carriers, we find several predicted to be related to GSH metabolic processes. Most notably, SLC25A39, the recently identified GSH transporter is amongst our top hits (ranked 4^th^). Furthermore, several SLC25 which are known to be relevant to mGSH metabolism are amongst the top candidates: SLC25A1^69,70^ (ranked 1^st^), SLC25A10^71,72^ (ranked 2^nd^), SLC25A13^73^ (ranked 3^rd^), and the iron transporter SLC25A37^16^ (ranked 8^th^). Surprisingly, the homolog of SLC25A39, SLC25A40, is not predicted to be involved in GSH metabolism, ranking 37^th^ of the 53 SLC25 family proteins. This may be due to the very low expression of SLC25A40 across the CCLE cell lines (**Supp. Fig.** 2), suggesting that SLC25A40 is quantitatively less relevant to mGSH transport in cancer cells. Other potential mitochondrial carriers of interest include SLC25A43 (ranked 6^th^), SLC25A24 (ranked 7^th^), and the OMM transporter SLC25A50 (ranked 5^th^). SLC25A43 is an orphan transporter which has recently been found to affect redox homeostasis^74^, but the other candidates have not been, to our knowledge, associated with GSH metabolism.

Structural similarities in the candidates, specifically within the tunnel region of the proteins, can inform potential roles of these transporters in mGSH metabolism. We investigated this through sequence-independent structural alignment of SLC25 transporters using TMalign. We find that, aside from its homolog SLC25A40, SLC25A39 shows the most similarities to our candidates SLC25A43, and SLC25A24. Important GSH binding residues in SLC25A39 are somewhat conserved in our candidates where, specifically, SLC25A43 shows a shift in residue position within the transporter tunnel and a substitution of an aspartate to a lysine residue, and SLC25A24 substitutes a threonine. These findings indicate that despite overall similarities in transporter tunnels between the GSH transporter and our candidates SLC25A43 and SLC25A24, substrate binding and transport mechanism are likely somewhat different, e.g., binding to alternate forms of GSH species such as GSSG or glutathione esters.

Expanding our search beyond the SLC25 family, 27 genes were classified with high probability (>0.80) for all classification tasks. Of these, some of the most probable candidates are discussed here for their possible roles in mGSH metabolism and transport.

Pyruvate carboxylase (PC) catalyzes the conversion of pyruvate to oxaloacetate in mitochondria. PC links glucose to GSH synthesis and plays roles in the control of redox and oxidative stress^75^. Recently, pyruvate metabolism has been associated with GSH metabolism and ferroptosis in lung cancer cells through the plasma membrane cysteine transporter SLC7A11^76^ (ranked 3^rd^ by GSH metabolic process GO term annotation probability in RF classifiers. PC is reported to specifically localize to the mitochondrial matrix with little evidence supporting it being membrane-bound; thus, it may be that PC interacts with the IMM-bound transporters to facilitate and modulate mGSH transport.

Mitochondrial calcium uniporter (MCU) is also a top hit in our models and a well-known mitochondrial transporter. Associated with its role in Ca^2+^ homeostasis are roles in iron and redox homeostasis, and cell death pathways^77^. With respect to GSH, MCU is regulated via s-glutathionylation^77^ and downregulation is associated with enriched GSH metabolism in melanoma cells^78^. It is possible that MCU plays a greater role in mitochondrial iron and GSH metabolism and transport in the context of cancer metabolism than has been reported thus far.

ATP-binding cassette family B6 (ABCB6) is a transporter protein with evidence for both plasma membrane and mitochondrial localization^79^. This protein transports the heme precursor porphyrin^80^, however the mechanism of transport has been shown to be GSH-dependent, with significantly increased activity in the presence of GSH^81^. These suggest GSH-mediated transport of substrates by ABCB6 but it is unclear if GSH merely modulates transport, or if it is transported as well, likely through GSH-conjugated species. Similarly, neudesin neurotrophic factor (NENF) is a mitochondrial localized protein primarily involved in the differentiation/development of neuronal cells^82^. NENF activity is modulated by the binding of heme^83^, which presents a connection to GSH that, to the best of our knowledge, has not yet been explored. Some evidence suggests NENF is membrane-associated^82^.

The results of our models find several genes that may play roles in mGSH metabolism and transport, and thus could guide future experimental analyses. Our findings are limited by our relatively small training dataset, in which more data would improve model performance and confidence of predictions. Furthermore, our models are generalized across cancer cell data from a range of tissue and cancer types. Future work, leveraging larger datasets, or datasets of specific cancer types would provide promising avenues for uncovering interactions of high relevance to the specific diseases. Another limitation of this work is the use of gene ontology terms for identifying relevant genes. Specific and manually curated databases for these functions, similar to MitoCarta, likely provide a more accurate set of relevant genes; however, these databases are limited. Using a database like gene ontology provides a resource for the annotation of many functions, processes, and cell localizations as demonstrated here, which cannot be replicated by specific databases.

Finally, the classifications from the individual models provide novel candidates for areas beyond the scope in this work. For example, the classifiers for other metabolites here were used for validation of the glutathione classifier, but predicted relevant genes for those metabolites were not investigated. Nevertheless, we anticipate that our findings here can be instrumental in the identification of potentially novel proteins involved in mitochondrial glutathione metabolism and transport in cancer cells. The results here and classifier code provide a tool to accelerate knowledge discovery and identify potential target genes for other biological functions and contexts.

## Abbreviations

*GSH*: Glutathione
*mGSH*: Mitochondrial glutathione
*ML*: Machine learning
*DL*: Deep learning
*ROS*: Reactive oxygen species
*OXPHOS*: Oxidative phosphorylation
*KEGG*: Kyoto Encyclopedia of Genes and Genomes
*CCLE*: Cancer Cell Line Encyclopedia
*GO*: Gene Ontology
*TrSSP*: The Transporter Substrate Specificity Prediction Server
*GSSG*: Glutathione disulfide
*PCA*: Principal component analysis
*PC*: Principal component
*RF*: Random Forest
*DT*: Decision Tree
*NB*: Naive Bayes
*SVM*: Support Vector Machine
*ROC*: Receiver operating characteristic
*PRC*: Precision-recall curve
*TP*: True positive
*TN*: True negative
*FP*: False positive
*FN*: False negative
*AUROC*: Area under receiver operating characteristic curve
*AUPRC*: Area under precision-recall curve
*MCC*: Matthew’s correlation coefficient
*CNN*: Convolutional neural network
*TPR*: True positive rate

## Ethics approval and consent to participate

Not applicable.

## Consent for publication

Not applicable.

## Availability of data and materials

All data and code used and generated in this work are publicly available at https://github.com/lkenn012/mGSH_cancerClassifiers^84^.

## Author contributions

L.K. designed and trained models and performed all other analyses. L.K. M.-E.H., and M.C.-C. interpreted the results and wrote the manuscript. M.-E.H., and M.C.-C., jointly supervised the study and acquired funding. L.K. M.-E.H., and M.C.-C. envisioned the study. All authors provided feedback and suggestions to develop the research and manuscript. All authors reviewed the manuscript.

