## Supplementary Figures for "A hybrid machine learning framework for functional annotation applied to mitochondrial glutathione metabolism and transport in cancers"

|  |  | **Metric** |  |
| --- | --- | --- | --- |
| Classifier model | MCC | AUROC | AUPRC |
| Glutamate metabolic process | 0.4835 | 0.8223 | 0.8110 |
| 2-oxoglutarate metabolic process | 0.5387 | 0.8224 | 0.8245 |
| Carnitine metabolic process | 0.4602 | 0.8107 | 0.8207 |

**Supplementary table 1: Model evaluations of non-GSH metabolite RF classifiers.** Values are mean metrics across feature sets.

|  | **RF classifier model** | | |
| --- | --- | --- | --- |
| Gene Symbol | GSH | Mitochondria | Transporter |
| GSR | 1 | 0.9223  (0.0131) | 0.8558  (0.0203) |
| ETHE1 | 1 | 0.8648  (0.0425) | 0.8312  (0.0254) |
| PARK7 | 1 | 1 | 0.8120  (0.0092) |
| GLRX2 | 1 | 1 | 0.8252  (0.0134) |
| ALDH2 | 0.8809  (0.0136) | 1 | 0.8383  (0.0284) |
| IDH1 | 0.8778  (0.0195) | 0.8900  (0.0080) | 0.8440  (0.0171) |
| NENF | 0.8754  (0.0122) | 1 | 0.8042  (0.0353) |
| PC | 0.8638  (0.0141) | 0.9265  (0.0083) | 0.9315  (0.0180) |
| ABCB6 | 0.8542  (0.0127) | 0.9109  (0.0046) | 1 |
| OAT | 0.8496  (0.0110) | 0.9547  (0.0087) | 0.8755  (0.0146) |
| CPS1 | 0.8446  (0.0119) | 0.8516  (0.0289) | 0.8552  (0.0309) |
| SUCLG2 | 0.8405  (0.0110) | 0.9346  (0.0183) | 0.8648  (0.0222) |
| ACADVL | 0.8398  (0.0073) | 0.9245  (0.0256) | 0.8523  (0.0318) |
| MCU | 0.8314  (0.0072) | 1 | 1 |
| TIMM9 | 0.8199  (0.0082) | 0.9537  (0.0115) | 0.8742  (0.0215) |

**Supplementary table** **2: Most probable non-SLC25 candidate mGSH transporters by mean RF classification probabilities with standard error, ranked by GSH probability**. Probability of 1 indicates the gene is already annotated by corresponding GO term.

|  |  | Classifier | |
| --- | --- | --- | --- |
| GO term | Classification threshold | DeepGOPlus | Random forest |
| GO:0006749 glutathione metabolic process | 0.4 | 0.053 | **0.850** |
|  | 0.5 | 0.026 | **0.800** |
|  | 0.6 | 0.000 | **0.650** |
| GO:0005739 mitochondrion | 0.4 | 0.507 | **0.932** |
|  | 0.5 | 0.320 | **0.887** |
|  | 0.6 | 0.033 | **0.777** |
| GO:0022857 transmembrane transporter activity | 0.4 | 0.727 | **0.966** |
|  | 0.5 | 0.593 | **0.894** |
|  | 0.6 | 0.387 | **0.771** |

**Supplementary table** **3**: **Sensitivity values for GO annotations by DeepGOPlus and RF (14 PC features) classifier models produced here**. Best performing models for each annotation task in bold.

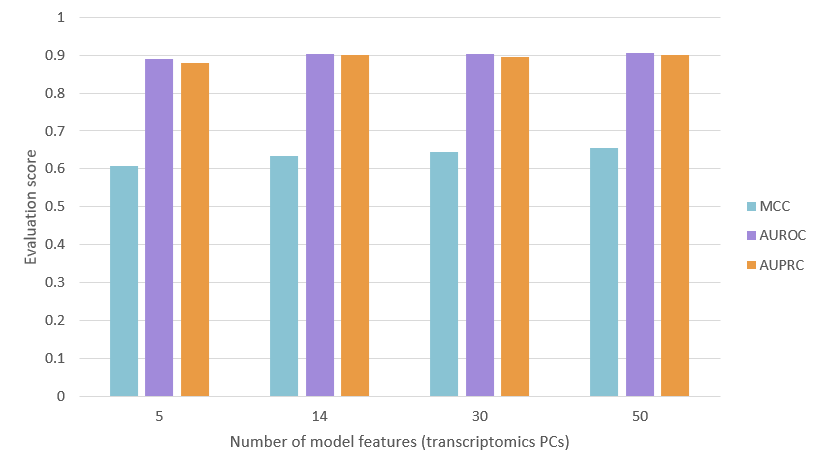

**Supplementary Figure 1:** Mean RF classifier evaluation metric scores with different numbers of features.

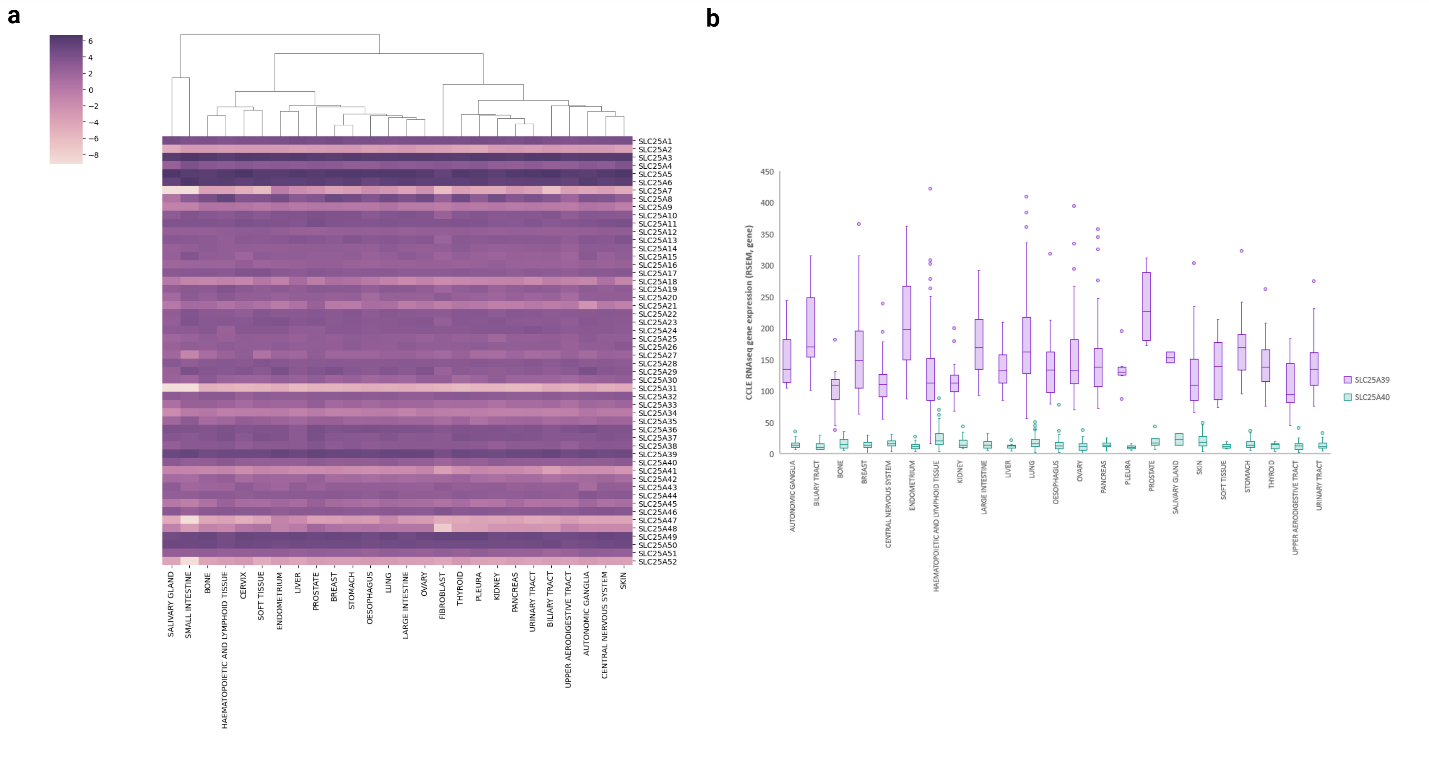
**Supplementary Figure 2:** CCLE transcriptomics of SLC25 family members. **a)** Mean SLC25 gene expression (log2 RNA-Seq by Expectation Maximization (RSEM)) across CCLE transcriptomics tissue types. Tissues are clustered using agglomerative clustering through ward linkages of euclidean distances.

**b)** RSEM values across CCLE tissue types for known GSH transporters SLC25A39 (purple) and SLC25A40 (green).

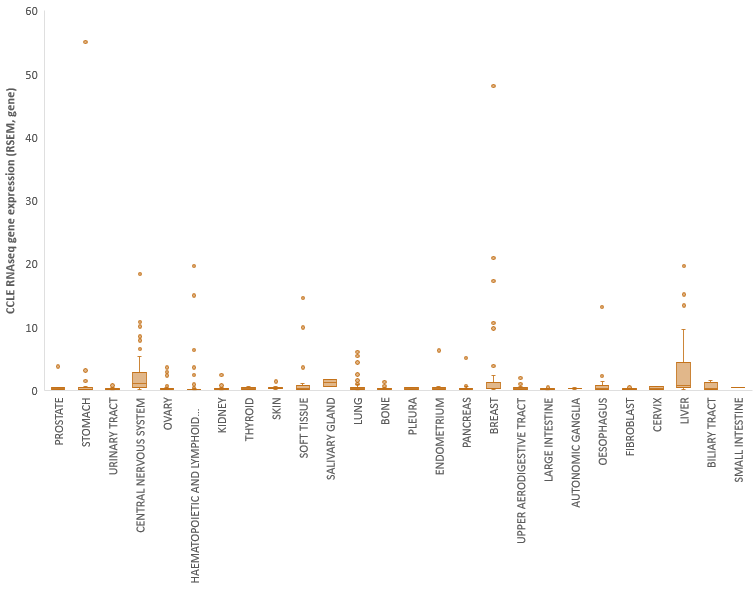

**Supplementary Figure 3:** CCLE transcriptomics profile of glutamate transporter SLC25A18. Expression values are RSEM values across CCLE tissue types.

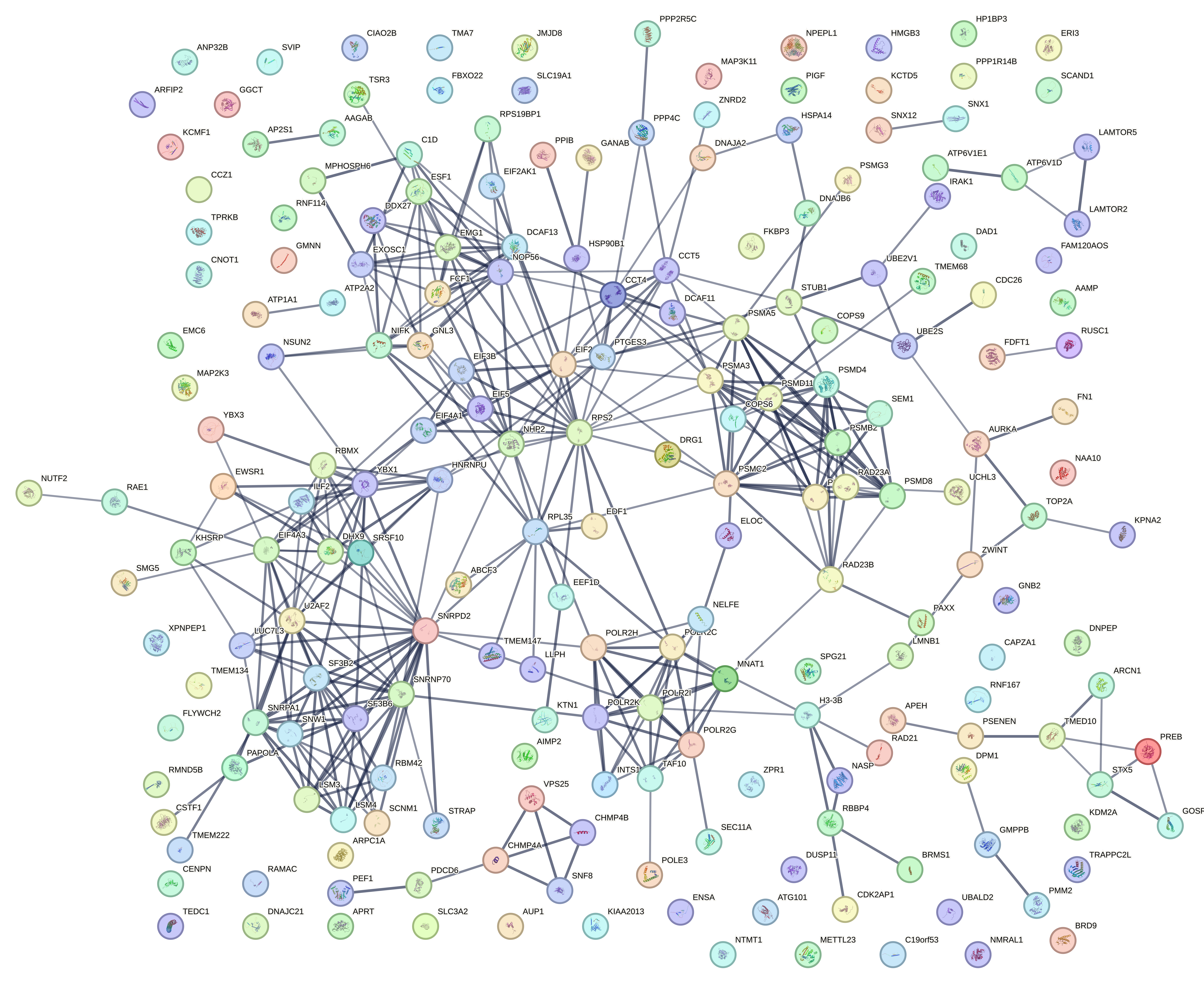
**Supplementary figure** **4**: STRING network of 206 genes predicted to localize to mitochondria by RF classifiers with no existing evidence in mitochondria databases. Strength of data support for connection is indicated by edge boldness (minimum confidence score of 0.70). 61 nodes with no edges, 86 nodes with no connection to the central network.

**A**

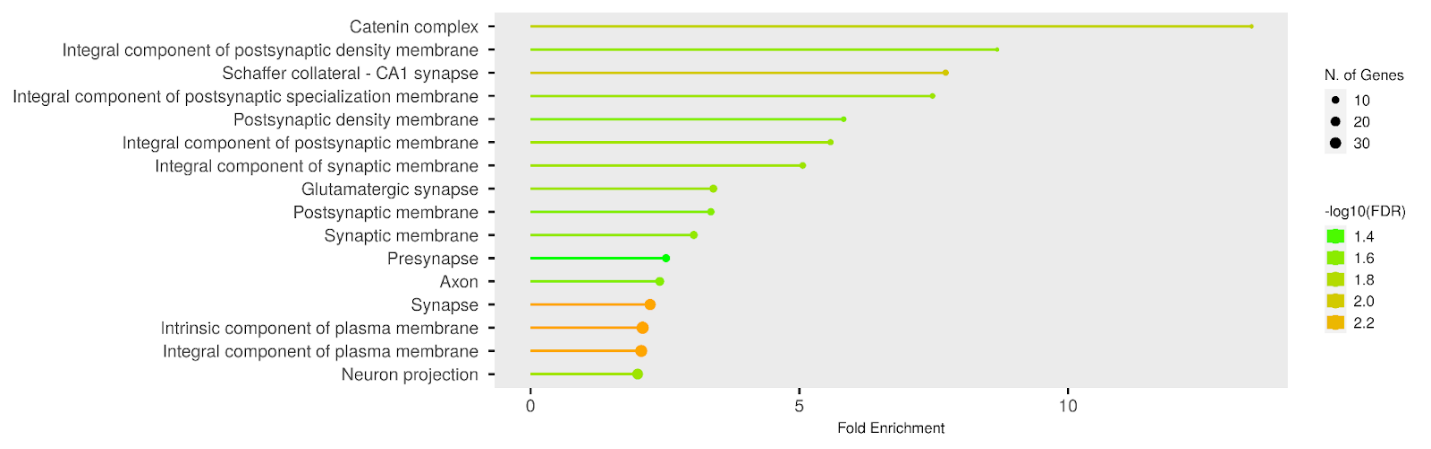
**B**
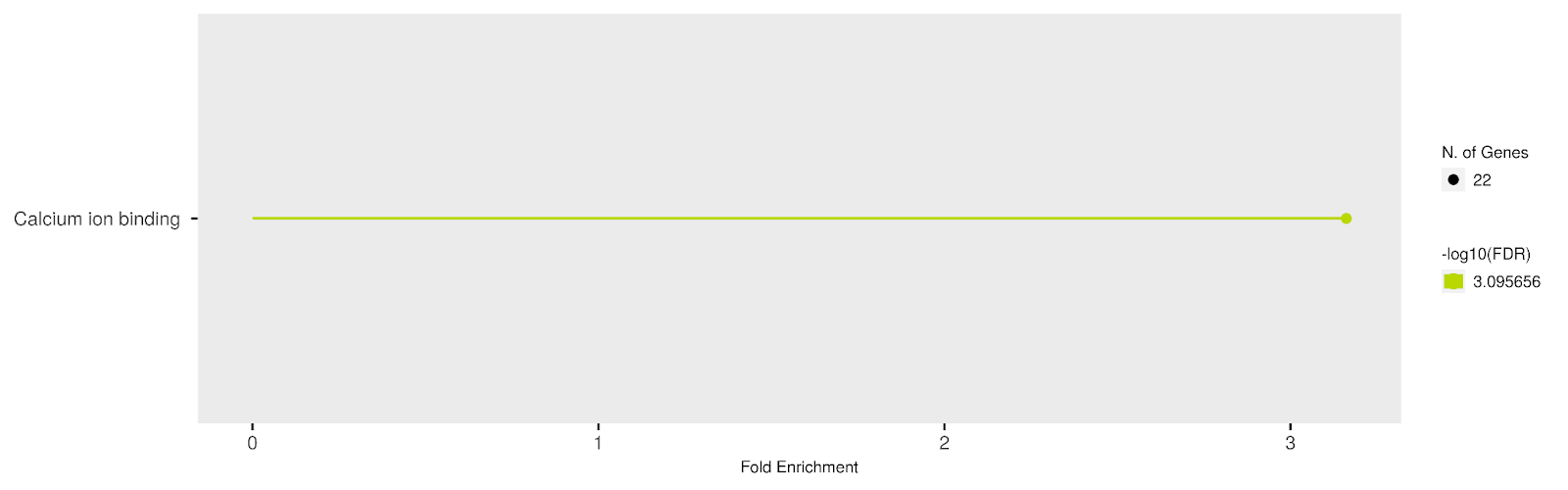
**C**
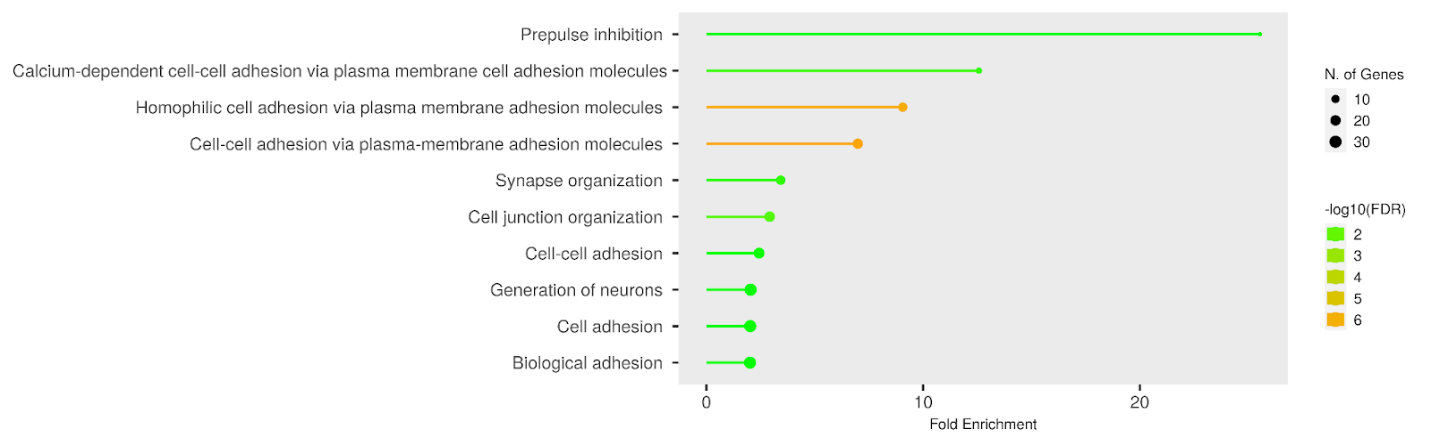

**Supplementary figure 5**. GO enrichment analysis of genes predicted to have transmembrane transporter activity by RF classifiers with no existing evidence in transporter databases. Enrichment plots of false positive transporter genes for cellular component (a), molecular function (b), and biological process (c), GO terms.

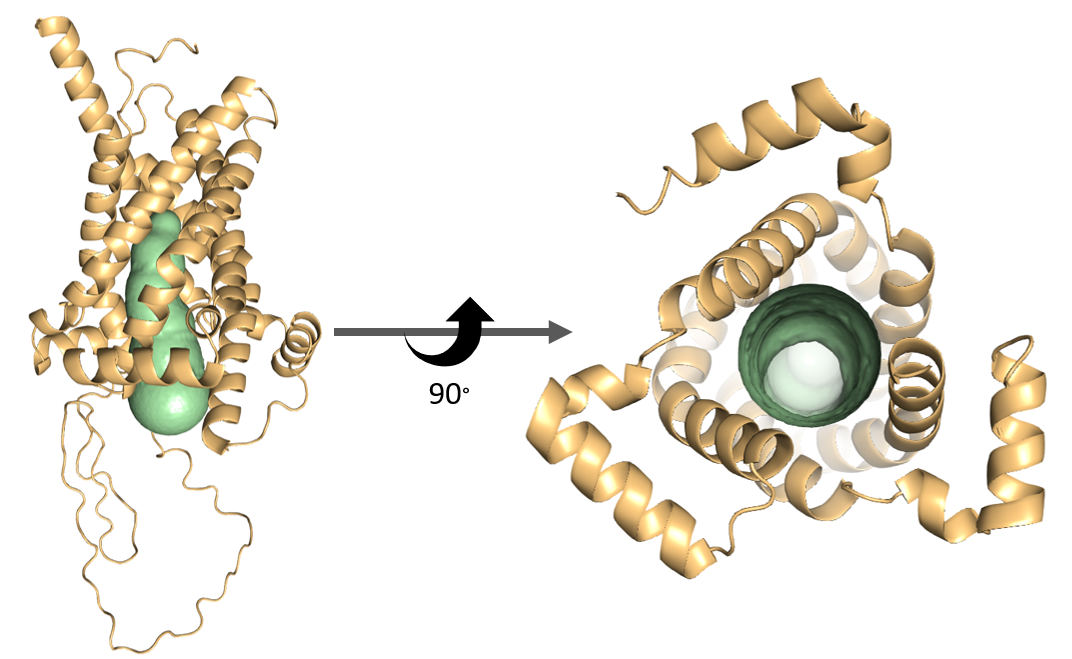

**Supplementary Figure 6:** CAVER identified transport tunnel residues in SLC25 structures. CAVER tunnel residue surface indicated in green, these residues are then used for local TM-alignments to identify similarities in substrate binding and transport function. Protein structure is AlphaFold-predicted SLC25A39 structure.

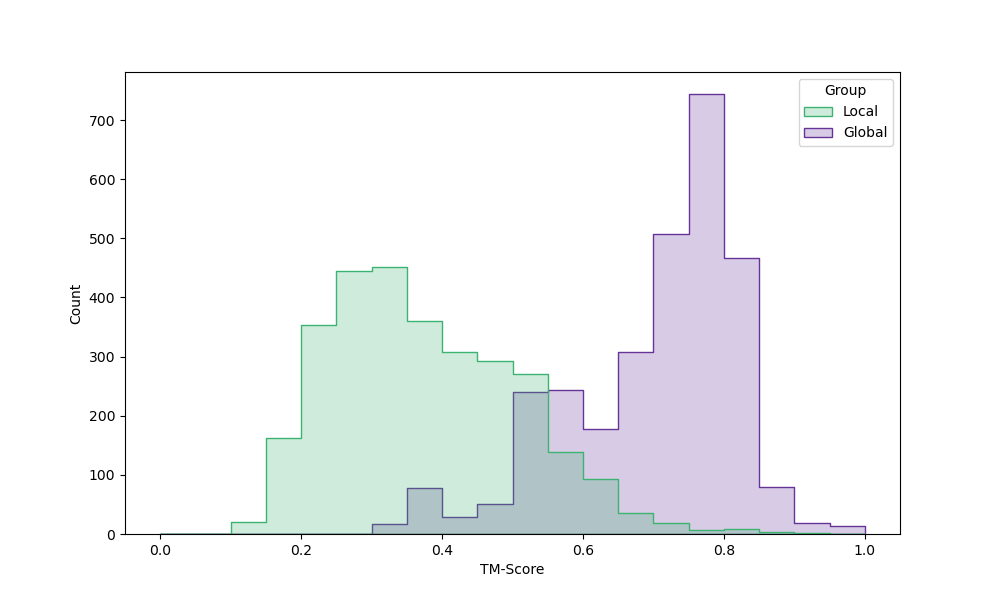

**Supplementary Figure 7**: Distribution of alignment scores by TM-align for pairs of SLC25 structures using entire protein structures (“Global”, purple) or tunnel residues only (“Local”, green) to calculate alignment.
